# Natural variation in temperature-resilient immunity in Arabidopsis

**DOI:** 10.64898/2026.08.17.745235

**Authors:** Richard Hilleary, Reza Sohrabi, Hannah McMillan, Saunia Withers, Jong Hum Kim, Sheng Yang He

## Abstract

Elevated temperature has been shown to compromise salicylic acid (SA)-mediated immunity in plants. The *Arabidopsis thaliana* accession C24 retains constitutively elevated SA and resistance to the hemibiotrophic pathogen *Pseudomonas syringae* pv. tomato DC3000 (*Pst* DC3000) at elevated temperature. C24 exhibits reduced biomass compared to that of a commonly studied accession, Col-0, in which SA-mediated immunity is compromised at elevated temperature. Neither the genetic basis of temperature-resilient immunity (TRI) nor the apparent growth-defense tradeoff in C24 is known. Here, we show that a Col-0 x C24 recombinant inbred line (RIL) population resolves TRI to a chromosome 5 locus accounting for most of the mapped genetic variance. This locus (named *TRI* hereinafter) coincides with a hotspot of structural rearrangement between the two accessions and includes a calcium-sensor gene (*CBL9*) and several NLR-type paralogs found only in C24. Consistent with a calcium-dependent signaling component, C24 mounts an elevated cytosolic Ca²⁺ response to *Pst* DC3000. Surprisingly, across the RIL population, disease resistance and biomass are only weakly correlated, with some lines exhibiting both large biomass and high pathogen resistance. These results show that temperature-resilient disease resistance is not only genetically tractable in C24 but also can be uncoupled from biomass cost. The *TRI* locus in C24 therefore encodes a natural mechanism(s) of temperature-resilient immunity with the growth-defense tradeoff resolved.

## Introduction

Rising global temperatures suppress yields of major food crops and are projected to cause substantial further losses in wheat, maize, rice, and soybean as warming continues (Lobell *et al*., 2011; Zhao *et al*., 2017). Compounding this direct yield penalty, pathogens and pests claim a large share of potential yield in these staple crops every year (Savary *et al*., 2019), and climate change may further destabilize plant-pathogen interactions by enhancing pathogen fitness and/or eroding the host immune responses that would otherwise hold infection in check (Velasquez *et al*., 2018).

A major route by which elevated temperature affects plant defense is suppression of the production of salicylic acid (SA), the hormone that coordinates resistance to biotrophic and hemibiotrophic pathogens (Liu *et al*., 2026). Transient temperature elevations (TTEs) of only a few degrees above optimal growth temperature are sufficient to compromise SA biosynthesis and signaling, and correspondingly resistance to bacterial infection and *NLR* gene-mediated responses in the commonly used accession Col-0 of the model plant *Arabidopsis thaliana* (Wang *et al*., 2009a; Huot *et al*., 2017). This thermosensitivity has been traced to either the NLR proteins in the resistant genetic background (Whitham *et al*., 1996; Zhu *et al*., 2010; Demont *et al*., 2025) or the transcriptional regulators CBP60g and SARD1, master regulators of the SA biosynthetic enzyme *ISOCHORISMATE SYNTHASE 1* (*ICS1*), in the susceptible genetic background (Wang *et al*., 2009b; Sun *et al*., 2015). In the genetically susceptible Arabidopsis plants, elevated temperatures reduce the recruitment of the components of the transcription machinery, including the formation of GUANYLATE-BINDING PROTEIN-LIKE 3 (GBPL3) defense-activated condensates, required for CBP60g expression, and restoring *CBP60g* expression using temperature-insensitive promoter is sufficient to rescue SA production and immunity at warm temperature without an obvious growth penalty (Kim *et al*., 2022). Interestingly, ectopic tomato *CBP60g* or *SARD1* expression using a heat shock-inducible promoter also enhances immunity at elevated temperature (Mukherjee *et al*., 2025). Consistent with the concept that disease resistance conferred by NLR receptors and receptor-like kinases (RLKs) generally requires SA-dependent amplification of effector-triggered immunity, boosting *CBP60g/SARD1* expression enhances ETI in Arabidopsis at elevated temperature (Kim *et al*., 2022).

Currently it is not clear if temperature-vulnerable NLRs or CBP60g/SARD1 are the principal points of temperature impact in plant immunity. Additional thermosensitive nodes converge on the same pathway, including CAMTA-mediated repression of *ICS1/CBP60g/SARD1* at warm ambient temperature (Kim *et al*., 2013), temperature-attenuated EDS1/PAD4 immune complexes (Carstens *et al*., 2014), cross-talk with the growth-promoting PIF4-COP1-DET1 thermomorphogenesis module via the SUMO E3 ligase SIZ1 (Gangappa & Kumar, 2018; Hammoudi *et al*., 2018) and the thermoresponsive immune transcription factor bHLH059 (Bruessow *et al*., 2021). A comprehensive understanding of the genetic and molecular bases of temperature-resilient immunity is therefore important both for understanding plant immunity and for informing future resistance breeding. Natural variation among plant germplasms offers an important route to this genetic basis. Unlike the reference accession Columbia-0 (Col-0), the Arabidopsis accession C24 maintains constitutively elevated basal SA and reactive oxygen species and shows enhanced, broad-spectrum stress resilience including heat and disease without an accompanying yield penalty despite significant biomass tradeoff (Bechtold *et al*., 2010, 2018). C24 also confers non-host-resistance to a range of non-adapted *Pseudomonas syringae* pathovars (Mishina & Zeier, 2007), and transcriptomic and metabolomic comparisons confirm that C24 mounts a markedly different, more constitutively defended response than Col-0 during *Pseudomonas* infection (Orf *et al*., 2022). Chromosome-level assembly of C24 alongside other accessions has revealed 13–17 Mb of rearranged sequence and thousands of genes affected by copy-number and structural variation relative to Col-0, with rearrangement hotspots enriched for biotic-stress genes (Jiao & Schneeberger, 2020), underscoring that C24 is not simply a SNP-divergent version of Col-0 but carries substantial independent genomic architecture relevant to defense.

During this study, we discovered that C24 possess temperature-resilient immunity (TRI) in contrast to Col-0. To elucidate the molecular basis of TRI, we made use of an established Col-0 × C24 recombinant inbred line (RIL) population (Törjék *et al*., 2006b), previously deployed to dissect the genetic basis of flowering time and water-use efficiency (Ferguson *et al*., 2019), biomass and metabolite QTL (Lisec *et al*., 2007), and epistatic contributions to heterosis (Kusterer *et al*., 2007). Here, we combine large-scale phenotyping of this population infected with *Pst* DC3000 with linkage-based QTL mapping (Broman *et al*., 2019) to identify a large-effect locus on chromosome 5 that underlies TRI in C24. We further find that C24, but not Col-0, mounts a distinct calcium influx upon challenge with *Pst* DC3000, pointing to an additional, C24-specific layer of pathogen perception beyond its known constitutive SA elevation. Critically, the resistance loci identified here can be recombined away from the growth costs associated with them in the C24 parent, demonstrating that temperature-resilient immunity and vigorous growth are not obligately linked and pointing toward a genetically tractable route for breeding temperature-resilient plants with robust disease resistance and large biomass in a warming climate.

## Results

### C24 exhibits temperature-resilient disease resistance and SA accumulation

C24 exhibits a constitutively elevated SA level relative to Col-0 (Mishina & Zeier, 2007; Bechtold *et al*., 2010), so we first asked whether this phenotype, and the associated resistance to *Pst* DC3000, persists at elevated temperature. Unlike Col-0, C24 showed no visible disease symptoms including necrosis or chlorosis during *Pst* DC3000 infection at either 23 °C or 28 °C (Fig. 1A). Bacterial growth was modestly, though not significantly, lower in C24 at 23 °C compared to Col-0; at 28 °C, however, Col-0 supported significantly higher bacterial titers than C24, indicating that C24 retains disease resistance at elevated temperature (Fig. 1B). We next asked whether *CBP60g* expression and SA biosynthesis were elevated in C24 at elevated temperature during *Pst* DC3000 infection. At 23 °C, basal and induced *CBP60g* expression, SA, and SAG levels were comparable between C24 and Col-0 (Fig. 1C,D,E). At 28 °C, however, *Pst* DC3000 significantly induced *CBP60g* expression, SA, and SAG accumulation only in C24 (Fig. 1C,D), suggesting that temperature-resilient *CBP60g* expression and SA and SAG accumulation correlate with sustained defense in C24 at elevated temperature.

**Figure 1.**
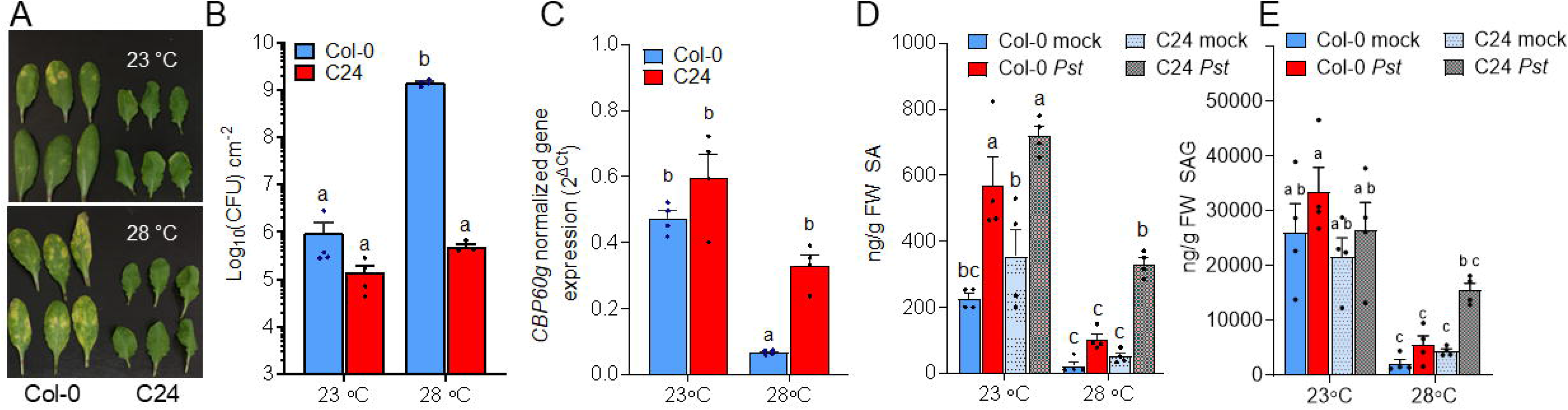
The C24 accession of Arabidopsis exhibits disease resistance resilience at elevated temperature. Col-0 and C24 plants were infiltrated with *Pst* DC3000 (1×10^6^ CFU ml^-1^). Images of infiltrated rosette leaves (A) and bacterial levels 3 dpi (B). *CBP60g* transcript (C), SA (D) and SAG (E) hormone levels at 1 dpi. Data in B-E are mean ± S.D. *n* = 3 (C) or 4 (B,D,E) biological replicates from one representative experiment analyzed with two-way ANOVA with Tukey’s honest significant difference (HSD) for significance. Experiments were independently performed three times, except for C, which was performed twice.

To test whether C24’s thermal disease resilience is inherited as a dominant trait, we assayed disease severity in Col-0 × C24 F_1_ hybrids at 23 °C and 28 °C (Fig. S1). F_1_ plants exhibited heterosis for leaf area at both temperatures and, surprisingly, showed markedly less disease symptoms than the Col-0 parent, suggesting enhanced defense in the F_1_ hybrids. Bacterial population quantification confirmed that *Pst* DC3000 growth in F_1_ hybrids was suppressed to levels resembling the C24 parent, consistent with dominant inheritance of C24’s resistance phenotype.

### Disease and SA phenotypes segregate transgressively within the Col-0 x C24 RIL population

To map genetic loci underlying disease resistance and SA accumulation, we assayed 443 Col-0 × C24 recombinant inbred lines (RILs), advanced by single-seed descent to the F_9_ generation and genotyped at 113 markers across the five chromosomes (Törjék *et al*., 2006a) for five traits following *Pst* DC3000 inoculation (by leaf infiltration): disease severity, *Pst* DC3000-induced SA and its glucoside conjugate SAG, and basal (mock-infiltrated with 0.25mM MgCl_2_) SA and SAG from the same lines. RILs were assayed in 11 sequential weekly batches (∼40–45 RILs per batch), with Col-0 and C24 included as parental controls in every batch.

Log_10_-transformed disease, SA (post infection with DC3000; DC3000 hereinafter), SAG (DC3000), SA (mock), and SAG (mock) were approximately continuous and unimodal across the RIL population, with standard deviations on the log10 scale ranging from 0.36 (SA, mock) to 0.86 (disease, DC3000) (Fig. 2; Table S1). For disease, SA (DC3000) and SAG (DC3000) RIL values extended beyond the parental range in both directions (Fig. 2), indicating transgressive segregation and a polygenic basis for these traits; SA (mock) and SAG (mock) instead shifted primarily below the C24 parental range, with fewer RILs exceeding it. Raw phenotype values also varied substantially by phenotyping batch independent of genotype, with batch means for disease varying up to ∼320-fold, and batch means for the SA/SAG traits varying 4- to 12-fold, across the 11 batches (Fig. S2). This variation was also present in the Col-0 and C24 controls included each week, which motivated including batch as an additive covariate in all QTL models.

**Figure 2.**
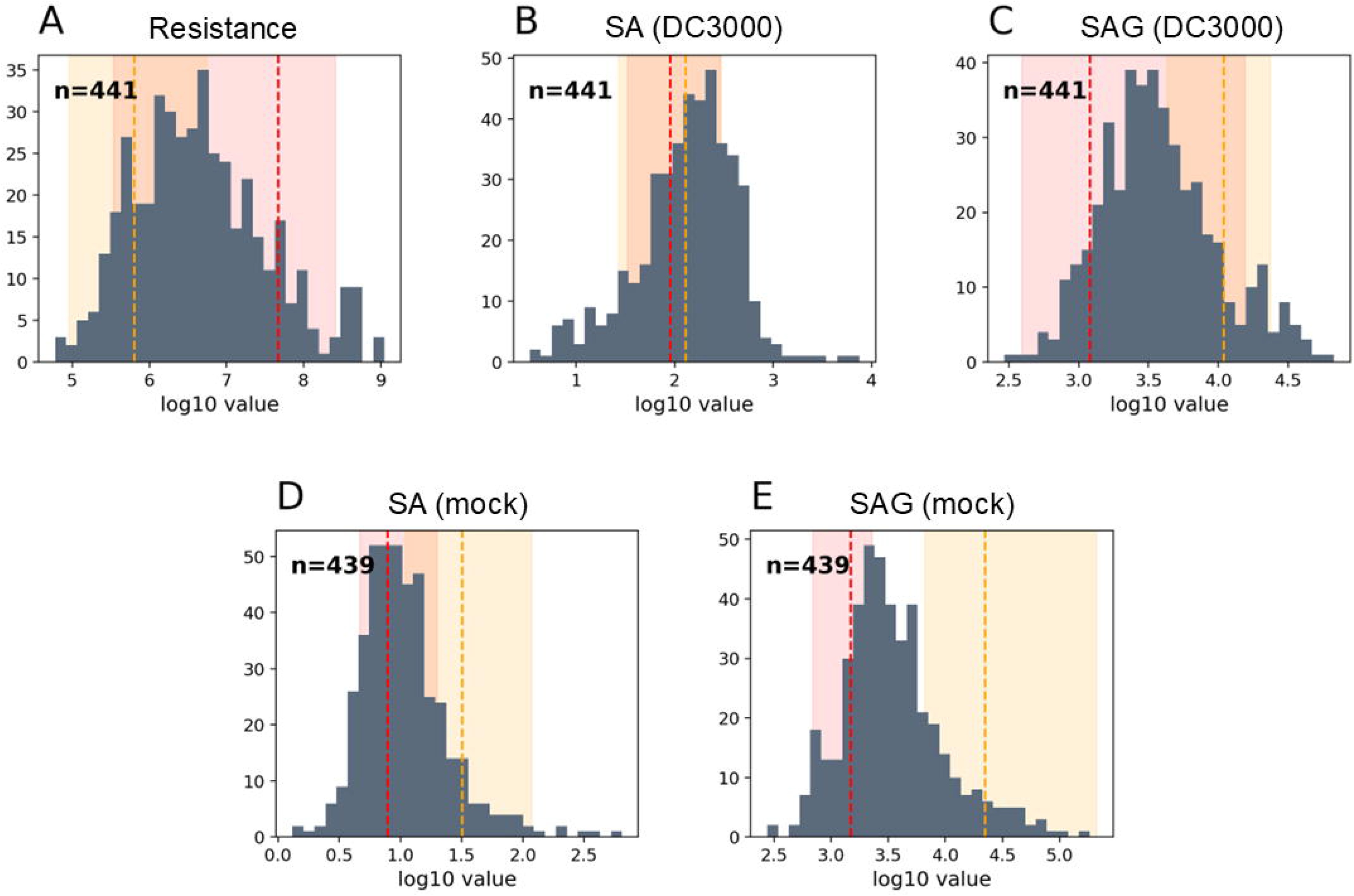
Phenotype distributions for traits in Col-0 x C24 RIL population. Histograms of log_10_-transformed disease resistance, SA (DC3000), SAG (DC3000), SA (mock), and SAG (mock) across all RIL phenotypes. Dashed vertical lines and shaded bands show the across-batch mean and across-batch mean range, respectively, of the Col-0 (red) and C24 (orange) parental controls for each trait. Sample sizes (n) reflect RILs with a valid measurement for that trait.

### QTL mapping identifies loci for disease resistance, SA, and SAG accumulation

Genome scans of the 441-RIL population based on phenotypes scores at 28°C identified a total of ten significant QTL peaks (permutation-based α = 0.05) across four of the five traits, with no genome-wide-significant locus detected for SA (mock) (Fig. 3, Table S1). Percent variance explained (PVE) ranged from 2.9% to 20.1% across these peaks, consistent with a mix of one large-effect and several moderate-effect loci rather than a single dominant locus. All loci are robust excluding a single flagged assay batch (batch 9; Methods), except for two nominal SA (mock) peaks, which do not survive batch 9 removal and are accordingly not reported as significant (Fig. S3).

**Figure 3.**
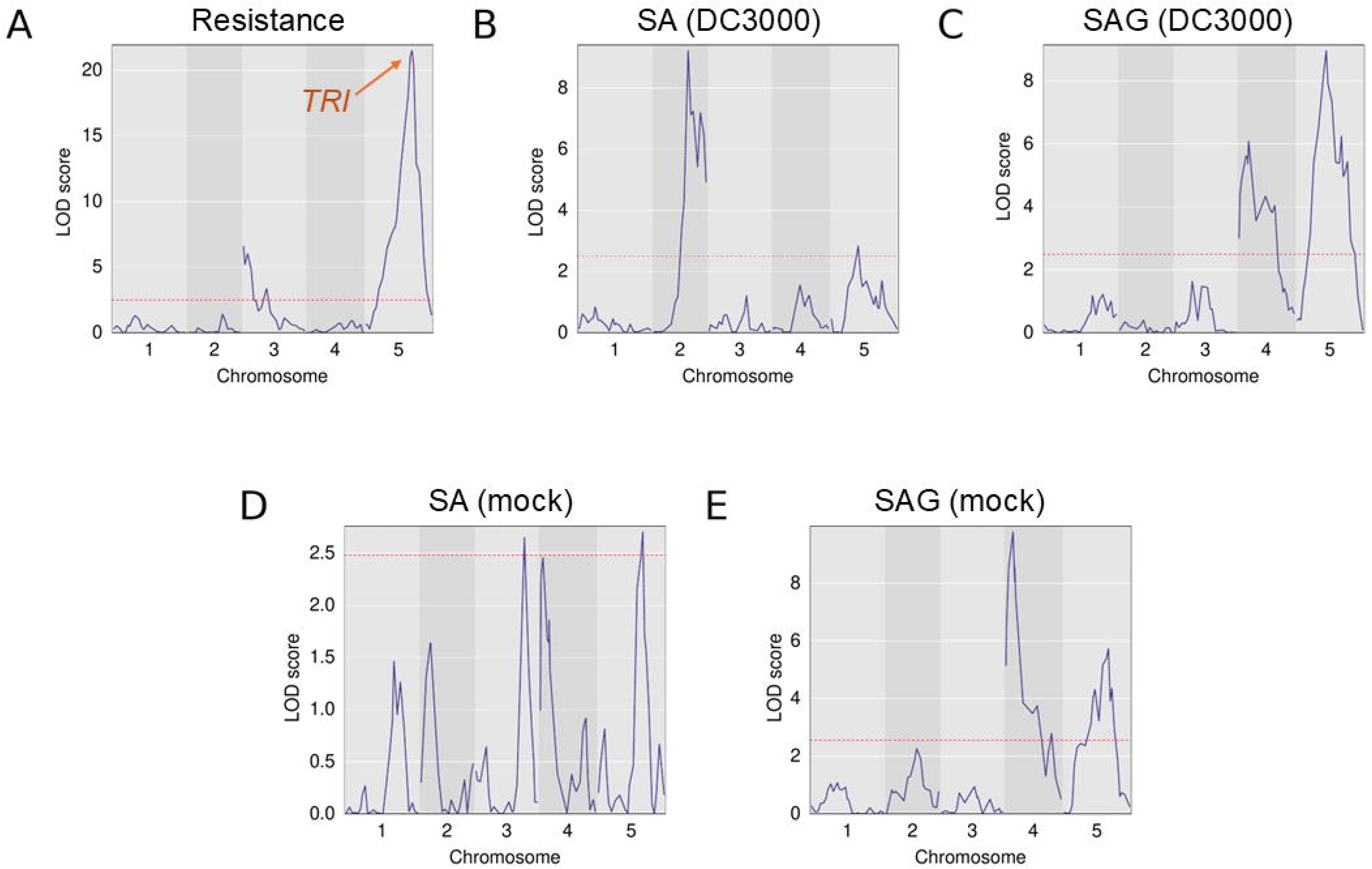
Genome-wide logarithm of odds (LOD) profiles for pathogen defense-associated traits. LOD profiles for (A) DC3000 resistance (*TRI* locus highlighted with orange arrow), (B) SA (DC3000), (C) SAG (DC3000), (D) SA (mock), and (E) SAG (mock) are presented. Dashed red line: permutation-based 5% significance threshold for that trait. Peak positions, LOD scores, 1.5-LOD support intervals, percentage variance explained (PVE), and the higher-value parental allele for each significant peak are reported in Table S1.

Disease severity was mapped to three loci, and at all three the Col-0 allele is associated with higher (more susceptible) bacterial titers. Two weaker loci lie near the top of chromosome 3, and a third, much stronger locus lies at 65.5 cM on chromosome 5 (LOD 21.50, PVE 20.1%, 1.5-LOD CI 60.1–71.6 cM). This locus is named *TRI*, for temperature resilient immunity, which by a wide margin was the strongest QTL in this dataset (Table S1). No genome-wide-significant locus was detected for basal SA. DC3000-induced SA mapped to two adjacent loci on chromosome 2, where the Col-0 allele was associated with higher induced SA (Table S1). A third DC3000-induced SA locus is located on chromosome 5 where the C24 allele is associated with higher induced SA (Table S1). The parental lines themselves do not differ significantly in DC3000-induced SA (mean SA across batches: Col-0 = 106 ng/g; C24 = 160 ng/g; t-test on log-transformed values, p = 0.24, n = 11 each), and the chr2 48.4cM locus is the single strongest QTL detected for the SA (DC3000) trait. RILs carrying the Col-0 allele at this locus average higher induced SA than either parent (317 ng/g, batch-adjusted 202 ng/g), while RILs carrying the C24 allele averaged near or below the Col-0 parent’s own value (185 ng/g, batch-adjusted 114 ng/g), which matches a pattern of transgressive segregation consistent with complementary gene action (Rieseberg *et al*., 1999). Both basal and DC3000-induced SAG mapped to chromosome 4 and chromosome 5 with the C24 allele associated with higher SAG (Table S1). Notably, the basal SAG QTL on chromosome 5 falls within the TRI locus. Taken together, resistance and SAG accumulation are both consistently linked to the C24 allele at every locus where they were detected, whereas induced SA shows the opposite pattern at its two largest-effect loci, indicating that C24’s resistance advantage tracks more closely with SAG accumulation than with free SA levels.

### Candidate genes in the TRI locus in association with disease resistance at elevated temperatures

Within the *TRI* locus, several candidate genes stand out (Table S2). Ranked by physical distance to the peak marker (Chr5:19,161,824), the closest gene of any kind is AT5G47229 (17.9 kb from the peak), an uncharacterized gene with no identifiable ortholog in the C24 assembly and lacking any functional annotation. The closest functionally annotated candidate is *CBL9* (AT5G47100, 31 kb from the peak), a calcium sensor of the calcineurin B-like family that negatively regulates defense responses, and *VICTR* and *VICTL* (AT5G46520/AT5G46510, ∼290–300 kb from the peak) are a tandem pair of TIR-NBS-LRR genes. *VICTR* encodes a functionally characterized EDS1/PAD4-dependent immune receptor that antagonizes abscisic acid signaling during *Pseudomonas* infection (Kim *et al*., 2012) and carries 90 coding SNPs plus a non-aligning segment relative to C24, the single most sequence-divergent gene identified anywhere in this interval, while its paralog *VICTL* carries a comparably large structural difference. The same tight cluster also includes *AT5G47260*, an uncharacterized NLR-family gene 27.6 kb from the peak, and its neighbor *ADR1-L3* (*AT5G47280*), 31.3 kb from the peak.

Between 573–575 kb from the peak, the interval also contains *CIPK19* and *CIPK20* (AT5G45810/AT5G45820), two more CBL-interacting protein kinases. Farther still, at 818 kb and 930 kb from the peak respectively, are *RLP56* (AT5G49290) and *NPR3* (AT5G45110). *RLP56*, an extracellular LRR receptor-like protein, carries 164 coding SNPs plus 13 indels and is the single most sequence-divergent gene identified anywhere in this interval. NPR3, a negative regulator of NPR1 turnover, carries 12 coding SNPs and a divergent promoter (17 SNPs, 8 indels).

Farther from the peak (830 kb–1 Mb), the interval also contains *RRS1B* and *RPS4B* (AT5G45050 and AT5G45060), a paralogous Toll/interleukin-1 receptor nucleotide-binding leucine-rich-repeat (TNL) gene pair approximately 60% identical to the canonical *RRS1/RPS4* pair, arising from a duplication that predates the Arabidopsis/Brassicaceae split (Saucet *et al*., 2015). *RRS1B* and *RPS4B* are both substantially more divergent in the C24 assembly than the canonical *RRS1/RPS4* pair, consistent with this locus being fast-evolving and duplication-prone. Consistent with this interval harboring multiple immune-gene candidates, whole-genome alignment between Col-0 and C24 shows the *TRI* locus to be substantially enriched for structural rearrangement relative to flanking sequence, with a dense concentration of highly diverged, duplicated, inverted, and Col-0-specific segments clustered around the peak-proximal candidates and the *RRS1B/RPS4B*-type cluster (Fig. 4). Mapping this same region onto the C24 assembly’s own coordinate system, identified 32 C24 genes with no Col-0 ortholog in the syntenic window, itself ∼800 kb–1 Mb from the peak in Col-0 coordinates. Four of these, clustered within a 200 kb span, are intact open reading frames carrying NB-ARC Walker-A/P-loop motifs found in plant NLR immune receptors. Two align to the canonical RPS4 and RRS1B proteins at near-full length and 49–51% identity (ATC24-5G63040 vs. RPS4, 1,121/1,149 aa aligned; ATC24-5G63050 vs. RRS1B, 605/754 aa aligned), consistent with paralog-level rather than background similarity. The other two show weaker, partial similarity to different NLR subclades and are not confidently assigned to a specific gene. The 1:1 C24 orthologs of Col-0 *RPS4/RRS1* (ATC24-5G63110/ATC24-5G63120) lie within the same cluster; C24 therefore retains the canonical gene pair alongside an additional, locally duplicated set of paralogs. A fifth, shorter (200 aa) intact ORF (ATC24-5G64570) carries no NB-ARC motif but is 65% identical to the NLR RPS6 over its N-terminal TIR-domain region and occupies RPS6’s syntenic position, indicating that the C24 copy of RPS6 may be truncated to a shortened TIR-domain remnant relative to the full-length Col-0 gene.

**Figure 4.**
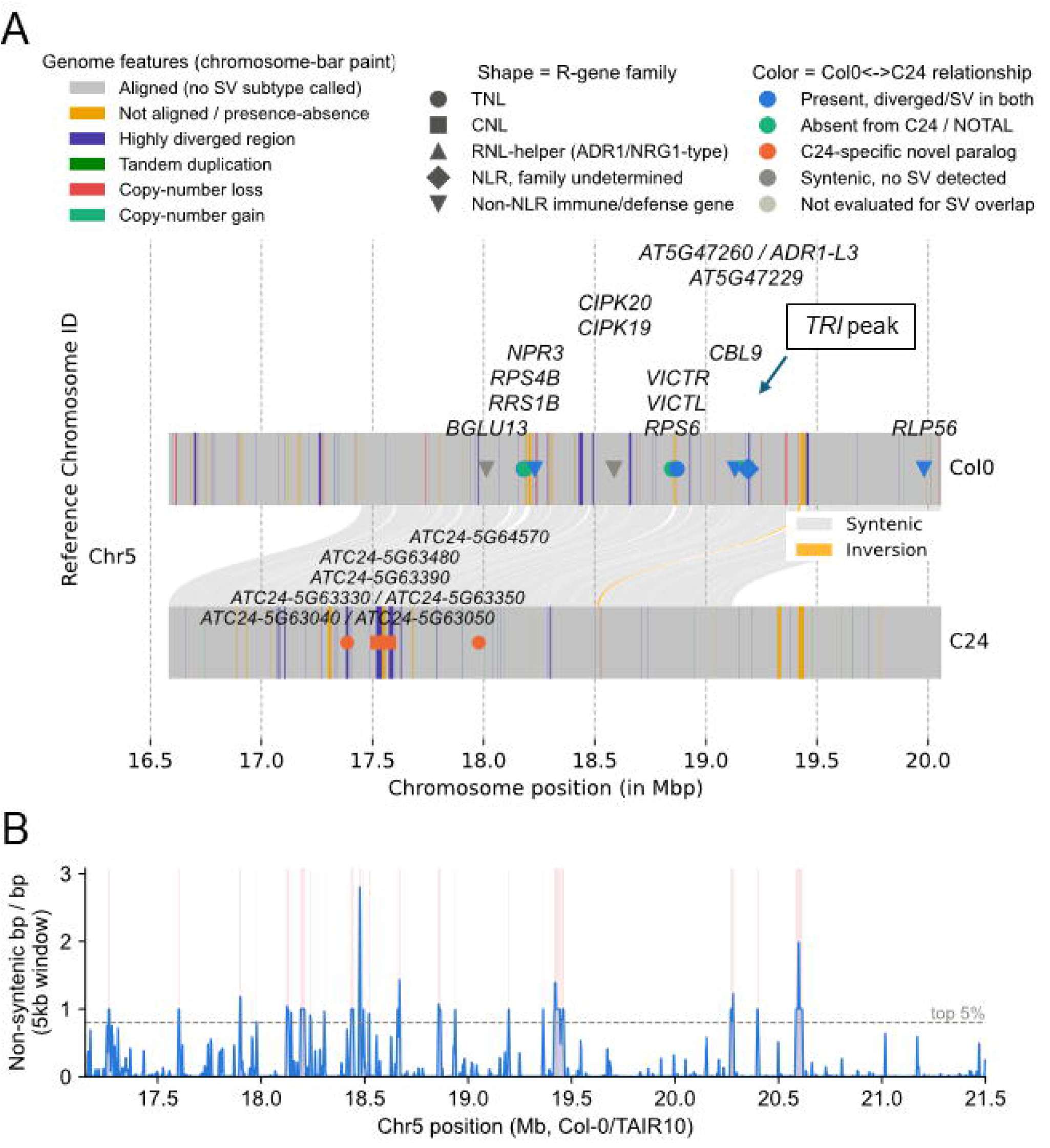
The *TRI* locus in chromosome 5. (A) Col-0×C24 synteny map across the QTL interval (Chr5:17,150,000–21,500,000, TAIR10). Ribbons connecting the Col-0 (top) and C24 (bottom) chromosome bars show synteny and rearrangement identifier (SyRI)’s top-level block classification (grey, syntenic; colored, inversion). Each chromosome bar is additionally painted with highly diverged regions, tandem duplications, local copy-number gain/loss, and non-aligning (presence/absence) segments. The 14 candidate genes (Table S2) are labeled at their Col-0 positions and, where an individual C24-specific paralog has been identified, at their C24 positions below. Marker shape indicates *R*-gene family and marker color indicates Col-0↔C24 relationship. Gray connecting lines link each C24-specific paralog cluster to its syntenic Col-0 window. (B) Local rearrangement-density score across the same interval. Summed base-pair coverage of all non-syntenic SyRI annotation in 5 kb sliding windows (1 kb step), following the window and step size of Jiao and Schneeberger (2020, Nat. Comm.). The grey dashed line marks the empirical top 5% of this density across the interval; pink-shaded windows meet or exceed that threshold and are candidate rearrangement hotspots.

This same chromosome 5 interval also contains QTLs for basal, uninfected SAG accumulation. SAG (mock) peaks on chromosome 5 at 63.0 cM (Chr5:18,282,539; LOD 5.74; PVE 5.8%), with the C24 allele associated with higher basal SAG and its 1.5-LOD support interval (41.7–71.6 cM) significantly overlaps with the *TRI* locus. Every candidate gene described above therefore also sits inside the SAG (mock) support interval. Whether this reflects a single pleiotropic locus or two distinct, tightly linked QTLs cannot be resolved at the current marker density (adjacent markers ∼0.9–1.4 Mb apart here) and would require finer-scale genotyping or a recombinant sub-population. The SAG (DC3000) QTL on chromosome 5 peaks at a separate, more distal position (41.4 cM, ∼11.1 Mb) that falls outside the *TRI* interval, indicating that SAG (mock) and SAG (DC3000) map as distinct QTL and that the *TRI* overlap is specific to basal, constitutive SAG accumulation rather than a pathogen-induced response.

Candidate genes were identified for the remaining mapped QTLs: two disease peaks on chromosome 3 (Dataset S3–S4), two SA (DC3000) peaks on chromosome 2 (Dataset S5–S6), the third SA (DC3000) peak on chromosome 5 (Dataset S7), the SAG (DC3000) loci on chromosomes 4 and 5 (Dataset S8–S9), and the SAG (mock) loci on chromosomes 4 and 5 (Dataset S10–S11). These intervals contain recognizable defense gene candidates, including *UGT74F1* and *UGT74F2*, the two Arabidopsis SA glucosyltransferases (George Thompson *et al*., 2017), *MPK6, MPK4, WRKY22/25/33*, several *CRK* and *CPK* gene clusters, the chitin receptor kinase gene *CERK1*, and *NDR1*.

### C24 exhibits robust calcium signaling during infection at elevated temperatures

Because one of the candidate genes at the *TRI* locus, *CBL9*, encodes a calcium sensor, we asked whether C24 also differs from Col-0 in pathogen-triggered calcium signaling itself by monitoring cytosolic Ca²⁺ dynamics in Col-0 and C24 rosette leaves expressing the R-GECO1-mTurquoise biosensor following infiltration with *Pst* DC3000 at 23°C and 28°C (Fig. 5A–C). C24 mounted a substantially larger Ca²⁺ response than Col-0 at both temperatures (AUC, Tukey’s p = 0.0085 at 23°C, p < 0.0001 at 28°C), indicating that C24 is poised for a robust response to a virulent pathogen at both standard and elevated temperatures.

**Figure 5.**
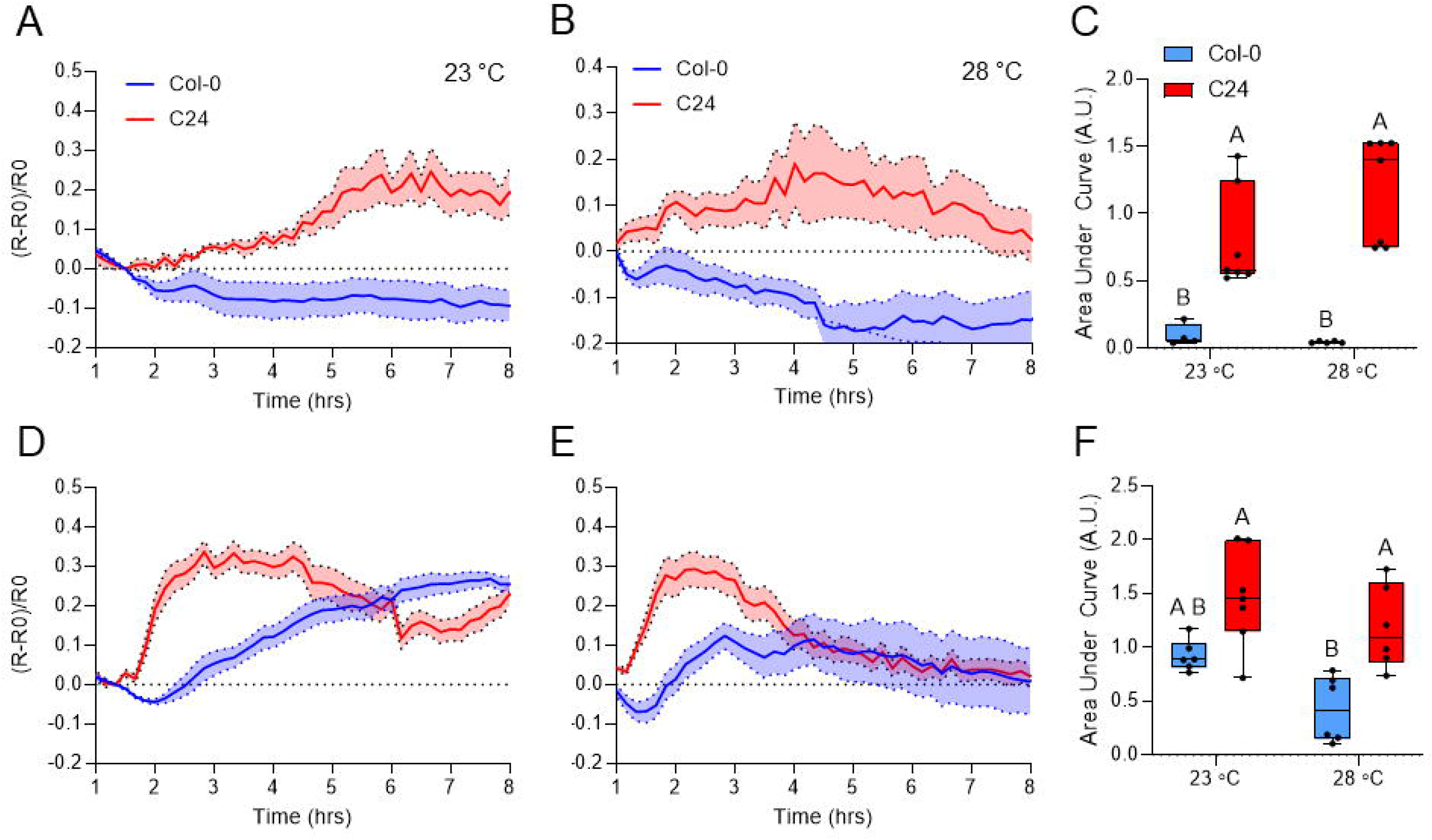
C24 exhibits strong, temperature-resilient calcium in response to pathogen challenge. Four-week-old Col-0 and C24 plants expressing the Ca²⁺ biosensor R-GECO1-mTurquoise were infiltrated with *Pst* DC3000 or *Pst* DC3000 (HopZ1a) and imaged at 23 °C or 28 °C. (A-C) Quantification of Ca²⁺ signals over time in rosette leaves infiltrated with *Pst* DC3000 at (A) 23 °C or (B) 28 °C, and (C) area under the curve values (AUC) observed over the time course. (D-F) Quantification of Ca²⁺ signals over time in rosette leaves infiltrated with *Pst* DC3000 (HopZ1a) at (D) 23 °C or (E) 28 °C, and (F) AUC values. Data are mean ± S.E.M. (n = 4 to 7 [A-C] or 6 to 7 [D-F]) from one representative experiment. Experiment was repeated twice with similar results. Two-way ANOVA with Tukey’s test for significance (p<0.05) was performed for peak maxima analyses with letters denoting differences in statistical significance.

*Pst* DC3000 delivering the effector HopZ1a triggers ZAR1/ZED1-dependent ETI in Arabidopsis (Lewis *et al*., 2013). We found that *Pst* DC3000 (HopZ1a) elicited a stronger Ca²⁺ response in both Col-0 and C24 at 23 °C compared *Pst* DC3000 alone, and only C24 mounted a robust Ca^2+^ response at 28 °C (Fig. 5D–F). Furthermore, C24 exhibited both a faster rise and an earlier peak in cytosolic Ca²⁺ signal compared to Col-0 at both temperatures in response to *Pst* DC3000 (HopZ1a) (Fig. S4). Time to half-maximal rise was shorter in C24 than Col-0 at 23 °C (2.0 ± 0.1 h vs. 2.9 ± 0.2 h) and, to a lesser extent, at 28 °C (1.8 ± 0.1 h vs. 2.1 ± 0.1 h), with peak time showing the same pattern (23 °C: 2.7 ± 0.2 h vs. 4.9 ± 0.4 h; 28 °C: 2.5 ± 0.3 h vs. 3.7 ± 0.4 h). Together, these results indicate that the enhanced Ca^2+^ response in C24 to pathogen challenge is distinguished not only by its magnitude but also by its speed, with C24 mounting a larger, faster, and earlier-peaking Ca²⁺ response than Col-0 across both virulent and ETI-eliciting pathogens and temperatures.

### Biomass and disease resistance segregate independently across the RIL population

Growth-defense tradeoffs are a longstanding obstacle to crop improvement, since resistance often comes at a cost to vegetative yield. Biomass in the Col-0 x C24 cross is known to vary substantially, including transgressive and heterotic effects exceeding both parents (Meyer *et al*., 2004; Kusterer *et al*., 2007; Lisec *et al*., 2007). However, it is not known if biomass and defense are obligately linked in the progeny of the Col-0 x C24 cross. To directly address this important question, we compared normalized biomass and normalized disease resistance (scaled to the Col-0/C24 parental range, scored at 28 °C) across 77 RILs with paired, quality-controlled measurements for both traits (Fig. S5). The two traits were only weakly and non-significantly correlated (r = -0.19, p = 0.10), indicating that enhanced growth does not come at an obligate cost to resistance across the population.

Both traits showed transgressive segregation, but to very different degrees. Z-score normalized biomass ranged from -0.32 to 3.72, with 38 of 77 RILs (49%) exceeding the parental range entirely; normalized resistance ranged from -0.69 to 2.46, with only 9 of 77 RILs (12%) at 28°C and 11 of 77 (14%) at 23 °C exceeding C24-level resistance. Disease resistance is therefore bounded much more tightly by the C24 parent than biomass is bounded by Col-0, consistent with resistance being governed by a small number of large-effect loci at which C24 already carries a near-optimal allele, while biomass remains free to vary polygenically.

This decoupling raises whether individual RILs can combine these traits in ways the parents cannot, and whether such combinations persist across temperature. Because our population screen scored resistance only at 28 °C, it could not test temperature-dependence directly. We therefore assayed two particular RILs, Q044 and Q017, at both 23 °C and 28 °C with additional replication (Fig. 6). Q044 and Q017 were chosen because, despite diverging substantially in biomass, both showed resistance exceeding Col-0 in preliminary screens. Specifically, biomass differed ∼3-fold between Q044 and Q017 (grams fresh weight: Q044 = 0.624 ± 0.024, Q017 = 0.187 ± 0.01), yet disease severity at 28 °C was indistinguishable between them (log_10_ CFU/cm²: Q044 7.30 ± 0.15; Q017 7.34 ± 0.25). Both RILs showed a smaller increase in temperature-induced disease susceptibility than Col-0 (Δlog_10_ CFU/cm², 23→28 °C: Col-0 +2.99; Q044 +1.28, p = 0.0029; Q017 +1.73, p = 0.023), indicating greater temperature resilience independent of biomass. These results indicate that disease resilience does not simply track plant size, supporting separable genetic control of biomass and pathogen defense.

**Figure 6.**
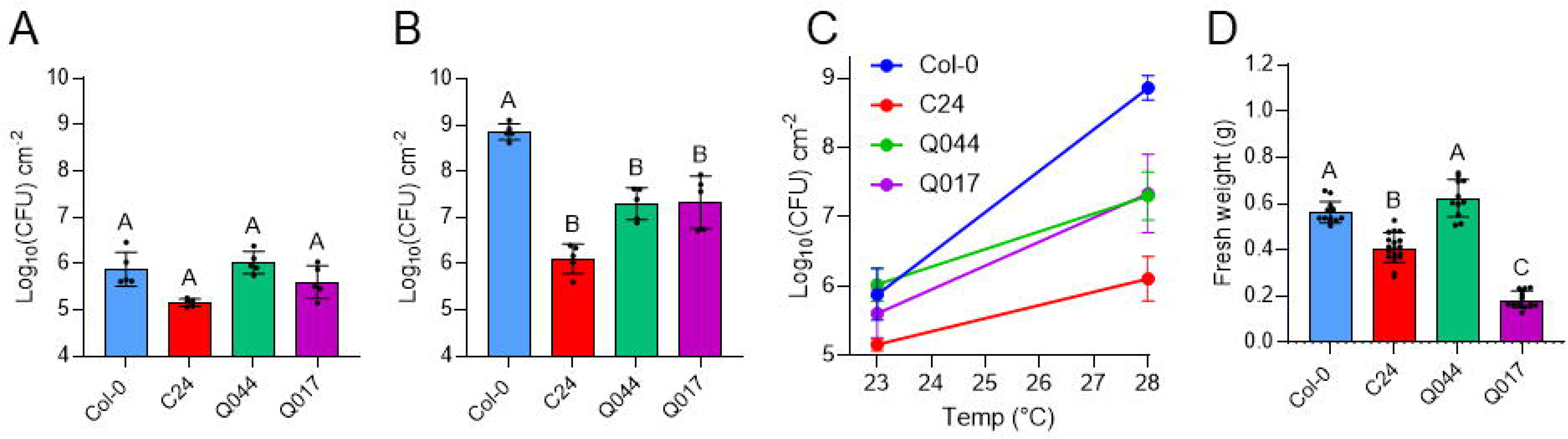
Temperature-resilient *Pst* DC3000 disease resistance can be decoupled from biomass. Disease severity and biomass for Col-0, C24, RILQ044, and RILQ017 at 23°C and 28°C. (A) 23°C disease severity (log_10_ CFU/cm²); mean ± SEM, (n = 5). (B) 28°C disease severity, mean ± SEM, (n = 5). (C) Mean log_10_ CFU/cm² at 23°C and 28°C per genotype, connected by line; slope reflects the temperature-induced susceptibility shift. (D) Biomass (fresh weight (g)), mean ± SEM, n = 11–18. Letter notation in A, B, D signifies significance (p < 0.05), one-way ANOVA, Tukey’s test for significance.

## Discussion

In this study, we show that temperature-resilient resistance in the natural Arabidopsis accession C24 resolves into a small number of genetic loci, with a single locus (*TRI*) on chromosome 5 accounting for most of the mapped genetic variance in disease resistance. Candidates at this locus span several distinct kinds of Col-0/C24 divergence rather than one clean category: extensive allelic divergence in genes present in both genomes (*CBL9, VICTR/VICTL, RRS1B/RPS4B*), a cluster of NLR-like paralogs found only in the C24 assembly with no corresponding Col-0 sequence, a shared gene (*RPS6*) apparently truncated to a TIR-domain remnant in C24, and, closest of all to the peak, an uncharacterized gene (AT5G47229) present in Col-0 but absent from the C24 assembly. C24 mounts a larger, faster, and earlier-peaking cytosolic Ca²⁺ response to pathogen challenge than Col-0 at both temperatures, showing that this heightened immune signaling persists as temperature rises. Despite a modest tradeoff between resistance and biomass at the *TRI* locus itself, growth and resistance segregate largely independently across the population, indicating that *TRI* can be recombined away from the growth cost associated with it in the C24 parent. Together, these findings establish *TRI* as a genetically rich, tractable natural locus for understanding climate-resilient immunity.

An unexpected finding from this study concerns the chromosome 2 locus for DC3000-induced SA (Table S1), where the Col-0 allele, not the C24 allele, is associated with higher induced SA, despite the parental lines themselves showing no significant difference in this trait (106 vs. 160 ng/g, p = 0.24, n = 11 each). The locus contains UGT74F1 and UGT74F2, the two Arabidopsis SA glucosyltransferases (George Thompson *et al*., 2017); because UGT74F2 preferentially conjugates SA to SGE rather than to SAG, and only SAG was measured here, differential UGT74F2 activity could lower free SA without a corresponding change in measured SAG, consistent with the absence of a SAG QTL at this locus in any of the four mapped SAG peaks. RILs carrying the Col-0 allele average higher induced SA than either parental line, a pattern of transgressive segregation possibly attributable to complementary gene action, where alleles of opposing effect, distributed across the two genomes, largely cancel in the parental lines themselves and are separable only upon recombination (Rieseberg *et al*., 1999; Gibson & Dworkin, 2004).

The gene nearest the chr5 disease-locus peak present in C24, *CBL9* (AT5G47100, 31 kb from the peak marker), encodes a calcineurin B-like calcium sensor previously implicated as a negative regulator of *Pseudomonas syringae* immunity. CBL9 and its paralog CBL1 activate CIPK6, which phosphorylates and suppresses RBOHD-mediated ROS production during PAMP- and effector-triggered immunity; *cbl1cbl9* and *cipk6* loss-of-function mutants both show enhanced resistance and SA accumulation upon *Pst* DC3000 challenge (Sardar *et al*., 2017; Vishwakarma *et al*., 2026).

Another important finding of this study is that growth and defense can be genetically uncoupled in the Col-0 x C24 RIL population. Specifically, biomass is controlled polygenically across many loci (Lisec *et al*., 2008), while resistance is concentrated at the *TRI* locus. During phenotyping we noted that several RILs combined dramatically larger biomass than the C24 parent with disease resistance that matched or exceeded it, even at elevated temperature; a striking result given the observed vegetative growth penalty of the C24 parent compared to the Col-0 parent. Results and RILs from this study therefore provide a good foundation for future molecular insights into decoupling growth from defense in a natural accession.

In summary, this work identifies a single, tractable chromosome 5 locus underlying temperature-resilient disease resistance in C24, nominates a short list of strong candidate genes, and shows that temperature-resilient immunity can be genetically separated from growth costs. As rising temperatures are expected to erode conventional SA-dependent immunity across model and crop systems, this locus and its candidate genes offer a promising target for achieving disease resistance without sacrificing biomass in a warming climate.

## Methods

### Plant materials and growth conditions

The Col-0 x C24 F_10_ RIL (Törjék *et al*., 2006b) population was kindly provided by Thomas Altmann’s lab. Plants were grown from vapor-phase sterilized seeds (Clough & Bent, 1998) stratified at 4 °C for 48 hours in 0.1% sterile agar. The seeds were pipetted directly onto potting soil (Arabidopsis Mix; 1:1:1 SureMix Perlite (Michigan Growers Product, Inc.) : medium vermiculite : coarse perlite) and domed to maintain high humidity for 1 week, after which domes were removed. Plants were grown in climate-controlled growth chambers with a 12hr diurnal cycle at 60% RH, 22 °C, and approximately 100 µmol m^-2^ s^-1^ white, fluorescent light. *UBQ10::R-GECO1-mTurquoise* C24 line was generated via floral dip (Bent, 2006) and T_3_ homozygous plants were used for experiments.

### Disease assays, hormone analysis, and biomass measurements

All disease assays were performed on four-week-old Arabidopsis plants by infiltrating a 10^6^ CFU mL^-1^ inoculum of *Pst* DC3000 suspended in 0.25mM MgCl_2_ into the rosette leaves with a needleless syringe. Leaf tissue for qPCR or hormone analysis was collected one day after inoculation (1 DPI) and flash frozen in liquid nitrogen. Bacterial populations were calculated three days after inoculation (3 DPI) by collecting 4mm leaf discs in sterile 0.25mM MgCl_2_ and plating 10uL of serial dilutions on Luria-Bertani rifampin (100 µg mL^-1^) plates to determine CFU cm^-2^. For temperature treatments, plants were placed into 23 °C or 28 °C growth chambers (12hr diurnal cycle at 60% RH, 22 °C, and approximately 100 µmol m^-2^ s^-1^ white, fluorescent light) for 48 hours prior to *Pst* DC3000 inoculation and placed back into these respective chambers after inoculation. Rossette fresh weight (FW) was measured by severing the stem with scissors and immediately recording the mass (g) on a digital scale.

### Quantitative PCR

100 milligrams of fresh leaf tissues were flash-frozen in liquid nitrogen and ground using a Geno/Grinder (Cole-Palmer). Plant RNA was extracted using a Qiagen Plant RNeasy Mini Kit following the manufacturer’s protocol, including on-column DNase I digestion. cDNA was synthesized by adding 100–300 ng of RNA to a solution of oligo-dT primers, dNTPs and M-MLV reverse transcriptase (Invitrogen). Approximately 1.5 ng of cDNA was mixed with the appropriate primers (sequences listed in Table S3) and SYBR master mix (Applied Biosystems). Quantitative PCR (qPCR) was run on a 7500 Fast Real-Time PCR system or QuantStudio 3 Real-Time PCR system (Applied Biosystems), with 2–4 biological replicates (and 3 technical replicates for each biological replicate) per experimental treatment. *PP2AA3* was used as an internal control to calcium 2^ΔCt^.

### Hormone profiling protocol

Plant hormones were extracted and quantified using a modified version of an established protocol from 1DPI leaf tissue (Huot *et al*., 2017). Methanol-based extractions were carried out with SA-d_4_ or SA-^13^C_6_ as internal standards. Extracts were filtered and analyzed on either an Acquity Ultra Performance Liquid Chromatography system with a Quattro Premier XE MS/MS (Waters) or a 1260 Infinity High Performance Liquid Chromatography system connected to a 6460 Triple Quadrupole mass spectrometer (Agilent). The column was maintained at 40 °C with a flow rate of 0.4 ml/min, and a gradient of mobile phases consisting of water + 0.1% formic acid (A) and methanol (B) was employed as follows: 0–0.5 min, 2% B; 0.5–3 min, 70% B; 3.5–4.5 min, 100% B; 4.51–6 min, 2% B, followed by 1 minute of equilibration. The analytes were then introduced into the Agilent jet stream electrospray ionization source in negative ion mode, with delta EMV (–) set at 200. The mass spectrometer source was configured with the following parameters: gas temperature at 300 °C, gas flow at 5 L/min, nebulizer at 45 psi, sheath gas temperature at 250 °C, sheath gas flow at 11 L/min, capillary voltage at 3,500 V, and nozzle voltage at 500 V. Multiple reaction monitoring (MRM) mode was used for data acquisition with these settings: dwell time at 50 ms, cell accelerator voltage at 4 V, fragmentor voltage at 90 V and collision energy at 16 V for SA and SA-d4; fragmentor voltage at 130 V. The following MRM transitions were monitored: SA (m/z 137→93), SA-d_4_ (m/z 141→97), and SAG (m/z 299.2 → 137.2). For peak selection and data integration, QuanLynx v4.1 software (Waters) or the Quantitative Analysis (for QQQ) program in MassHunter software (Agilent) was utilized. Hormone levels were calculated as previously described(Huot *et al*., 2017).

#### Whole plant Ca^2+^ imaging and analysis

Soil-grown *UBQ10::R-GECO1-mTurquoise* Arabidopsis (4 weeks old) were imaged in a custom Alligator Luminescence System (Cairn). The chamber was equipped with an iXon 888 EMCCD camera (Oxford Instruments, Andor Technology), and a custom Optospin emission filter switching unit (Cairn). A pE-4000 (CoolLED) unit was connected to a fiberoptic cable with 4-splitter terminal mounted leads for excitation illumination of plant samples. The pE-4000 and Optospin units were equipped with 25mm diameter mounted optical filters (Chroma Technology) for imaging mTurquoise (excitation = 435nm; excitation filter:ET440/40x; emission filter:ET480/30m)) and R-GECO1 (excitation = 550nm; excitation filter:ET560/25x; emission filter:ET620nm/60m. Image acquisition was controlled via ImageJ using µManager2.0 software. Image analysis was performed using FIJI. Briefly, image stacks were auto-thresholded using the Default settings in the Thresholder tool to set background pixel values to NaN, then the R-GECO1 channel was divided by the mTurquoise channel to obtain ratiometric images. The polygon tool was used to manually define region of interests (ROI)s around rosette leaves that were infiltrated with bacteria. All data presented was normalized using the expression (R-R0)/R0, where R is the average R-GECO1:mTurquoise ratio value for a ROI at a specific time point, and R0 is the initial ratio value for the same ROI

### QTL analysis

The 443 Col-0 × C24 RILs were genotyped at 113 markers spanning the five chromosomes using the map previously constructed (Törjék *et al*., 2006a). Marker order and spacing followed this published map without modification.

For phenotype quality control, disease severity, SA (DC3000), SAG (DC3000), SA (mock), and SAG (mock) were log10-transformed prior to analysis to stabilize variance across the ∼2–3 orders of magnitude spanned by each trait. RILs missing a valid measurement for a given trait (due to failed infiltration, insufficient tissue, or assay failure) were excluded from that trait’s scan only, resulting in the per-trait sample sizes shown in Fig. 2 (n = 441 for disease, SA (DC3000), and SAG (DC3000); n = 439 for SA (mock) and SAG (mock)) out of the 443 genotyped RILs. Batch (representing the week phenotyping was performed) was included as an additive covariate in every model below to account for the substantial batch-to-batch variation observed even in the Col-0 and C24 controls (Fig. S2).

For QTL mapping, genome scans were performed in R/qtl2 (Broman *et al*., 2019) using Haley-Knott regression on a riself cross object, with batch as an additive covariate. Genome-wide significance thresholds were established independently for each trait by permutation (1000 permutations, α = 0.05); thresholds ranged from approximately 2.4–2.9 LOD across traits (Table S1). Peaks were identified with find_peaks(), and 1.5-LOD support intervals were calculated for each significant peak. Percent variance explained (PVE) was approximated from the peak LOD score as 1 − 10^(−2LOD/n)^, where n is the number of phenotyped RILs for that trait.

One phenotyping batch (week 9) was identified as a candidate outlier because its Col-0 control showed anomalously low bacterial titers (log_10_ 5.5 CFU/cm²) relative to all other batches (log_10_ 7.5–8.4), consistent with a technical failure (*e.g.*, infiltration or inoculum problem). All scans were repeated without batch 9; peak positions, LOD scores, and thresholds from this slightly reduced dataset are reported alongside the full-dataset results (Table S1). All ten significant peaks were robust to the exclusion of batch 9 except two nominal SA (mock) peaks, which fell below threshold once batch 9 was removed and are accordingly not reported as genome-wide significant.

### Gene structural variant analysis landscape

The physical interval underlying the chromosome 5 disease-resistance QTL (60.1–71.6 cM, 1.5-LOD support interval) was defined by converting genetic to physical position using flanking and peak markers with known physical coordinates (V.103a, 60.092 cM/17,188,344 bp; V.105, 65.461 cM/19,161,824 bp; V.107, 71.578 cM/21,434,621 bp), with physical position interpolated linearly between anchors based on nearest-edge distance of genes. All Araport11-annotated genes within this interval were extracted from the TAIR10 reference.

Gene structural variation between Col-0 and C24 across this interval was assessed from a whole-genome SyRI alignment (Goel *et al*., 2019) of the C24 chromosome-scale assembly (Jiao & Schneeberger, 2020) to TAIR10, classifying each aligned block as syntenic, rearranged (inversion, translocation, duplication, or copy-number variant), or absent. Orthologs between Col-0 and C24 genes was assigned by OrthoFinder (Emms & Kelly, 2019). Col-0 genes with no C24 sequence in their orthogroup were classified as lacking a C24 ortholog. Four C24-specific NB-ARC-domain genes with no Col-0 ortholog and no direct syntenic counterpart were placed at interpolated positions using their nearest flanking syntenic (SYN-block) anchors on both assemblies.

### Biomass and disease resistance decoupling screen

To directly compare biomass and disease resistance within the same plants, a separate cohort of RILs was screened by dip inoculation. Four- to five-week-old plants were dip-inoculated in a 10^7^ CFU mL^-1^ suspension of *Pst* DC3000 and maintained at 23 °C or 28 °C. Bacterial titers were determined at 3 days post-inoculation (DPI), and rosette fresh weight (FW) was recorded from the same plants. Col-0 and C24 parental controls were included each week and used to normalize biomass and disease values across weeks. Biomass and 28 °C disease severity (log10 CFU/cm²) were normalized to a common scale using the parental means from each assay, where the normalized value = 2 × (raw value − C24 mean) / (Col-0 mean − C24 mean) − 1, such that C24 = −1 and Col-0 = +1 on both axes. Because Col-0 exhibited higher bacterial titers than C24, the normalized disease axis represents a susceptibility index, with higher values indicating greater susceptibility.

## RESOURCE AVAILABILITY

### Lead contacts

Further information or requests for resources and reagents should be directed to and will be fulfilled by the lead contacts, Richard Hilleary and Sheng Yang He.

### Material availability

All novel materials described in this paper will be made available upon request, subject to completion of an MTA.

### Data and code availability

All data reported in this paper will be shared by the lead contacts upon request.

Original code generated for this paper is available at https://github.com/DickHilleary/RILanalyses.

Additional information required to reanalyze the data reported in this paper is available from the lead contacts upon request.

## Acknowledgments

This study was supported by funding from funds from United States Department of Agriculture Postdoctoral Fellowship 2020-10954 to R.H. and Duke University to S.Y.H. S.Y.H. is an Investigator at Howard Hughes Medical Institute. We thank Thomas Altmann and his lab for creating and providing the RIL lines used in this study. We thank the excellent staff at Duke University Phytotron and Greenhouse for plant growth support.

## AUTHOR CONTRIBUTIONS

R.H. conceptualized and designed this study; R.H., R.S., J.K., H.M., and S.W. performed experiments; R.H. and S.Y.H. analyzed data; R.H. and S.Y.H. wrote the paper and all authors approved the final article.

## References

Bechtold U, Ferguson JN, Mullineaux PM. 2018. To defend or to grow: Lessons from Arabidopsis C24. Journal of Experimental Botany 69: 2809–2821.

Bechtold U, Lawson T, Mejia-Carranza J, Meyer RC, Brown IR, Altmann T, Ton J, Mullineaux PM. 2010. Constitutive salicylic acid defences do not compromise seed yield, drought tolerance and water productivity in the Arabidopsis accession C24. Plant, Cell & Environment 33: 1959–1973.

Bent A. 2006. Arabidopsis thaliana Floral Dip Transformation Method. In: Wang K, ed. Agrobacterium Protocols. Totowa, NJ: Humana Press, 87–104.

Broman KW, Gatti DM, Simecek P, Furlotte NA, Prins P, Sen Ś, Yandell BS, Churchill GA. 2019. R/qtl2: Software for mapping quantitative trait loci with high-dimensional data and multiparent populations. Genetics 211: 495–502.

Bruessow F, Bautor J, Hoffmann G, Yildiz I, Zeier J, Parker JE. 2021. Natural variation in temperature-modulated immunity uncovers transcription factor bHLH059 as a thermoresponsive regulator in Arabidopsis thaliana. PLOS Genetics 17: e1009290.

Carstens M, McCrindle TK, Adams N, Diener A, Guzha DT, Murray SL, Parker JE, Denby KJ, Ingle RA. 2014. Increased resistance to biotrophic pathogens in the Arabidopsis constitutive induced resistance 1 mutant is EDS1 and PAD4-dependent and modulated by environmental temperature. PLoS ONE 9: e109853.

Clough SJ, Bent AF. 1998. Floral dip: a simplified method for Agrobacterium -mediated transformation of Arabidopsis thaliana. The Plant Journal 16: 735–743.

Demont H, Remblière C, Culerrier R, Sauvaget M, Deslandes L, Bernoux M. 2025. Downstream signaling induced by several plant Toll/interleukin-1 receptor-containing immune proteins is stable at elevated temperature. Cell Reports 44.

Emms DM, Kelly S. 2019. OrthoFinder: phylogenetic orthology inference for comparative genomics. Genome Biology 20: 238.

Ferguson JN, Meyer RC, Edwards KD, Humphry M, Brendel O, Bechtold U. 2019. Accelerated flowering time reduces lifetime water use without penalizing reproductive performance in Arabidopsis. Plant, Cell & Environment 42: 1847–1867.

Gangappa SN, Kumar SV. 2018. DET1 and COP1 modulate the coordination of growth and immunity in response to key seasonal signals in Arabidopsis. Cell Reports 25: 29–37.e3.

George Thompson AM, Iancu CV, Neet KE, Dean JV, Choe J. 2017. Differences in salicylic acid glucose conjugations by UGT74F1 and UGT74F2 from Arabidopsis thaliana. Scientific Reports 7: 46629.

Gibson G, Dworkin I. 2004. Uncovering cryptic genetic variation. Nature Reviews Genetics 5: 681–690.

Goel M, Sun H, Jiao W-B, Schneeberger K. 2019. SyRI: finding genomic rearrangements and local sequence differences from whole-genome assemblies. Genome Biology 20: 277.

Hammoudi V, Fokkens L, Beerens B, Vlachakis G, Chatterjee S, Arroyo-Mateos M, Wackers PFK, Jonker MJ, van den Burg HA. 2018. The Arabidopsis SUMO E3 ligase SIZ1 mediates the temperature dependent trade-off between plant immunity and growth. PLoS Genetics 14: e1007157.

Huot B, Castroverde CDM, Velásquez AC, Hubbard E, Pulman JA, Yao J, Childs KL, Tsuda K, Montgomery BL, He SY. 2017. Dual impact of elevated temperature on plant defence and bacterial virulence in Arabidopsis. Nature Communications 8: 1–11.

Jiao W, Schneeberger K. 2020. Chromosome-level assemblies of multiple Arabidopsis genomes reveal hotspots of rearrangements with altered evolutionary dynamics. Nature Communications 11: 1–10.

Kim JH, Castroverde CDM, Huang S, Li C, Hilleary R, Seroka A, Sohrabi R, Medina-Yerena D, Huot B, Wang J, et al. 2022. Increasing the resilience of plant immunity to a warming climate. Nature 607: 339–344.

Kim T-H, Kunz H-H, Bhattacharjee S, Hauser F, Park J, Engineer C, Liu A, Ha T, Parker JE, Gassmann W, et al. 2012. Natural Variation in Small Molecule–Induced TIR-NB-LRR Signaling Induces Root Growth Arrest via EDS1- and PAD4-Complexed R Protein VICTR in Arabidopsis. The Plant Cell 24: 5177–5192.

Kim Y, Park S, Gilmour SJ, Thomashow MF. 2013. Roles of CAMTA transcription factors and salicylic acid in configuring the low-temperature transcriptome and freezing tolerance of Arabidopsis. The Plant Journal 75: 364–376.

Kusterer B, Muminovic J, Utz HF, Piepho HP, Barth S, Heckenberger M, Meyer RC, Altmann T, Melchinger AE. 2007. Analysis of a triple testcross design with recombinant inbred lines reveals a significant role of epistasis in heterosis for biomass-related traits in Arabidopsis. Genetics 175: 2009–2017.

Lewis JD, Lee AH-Y, Hassan JA, Wan J, Hurley B, Jhingree JR, Wang PW, Lo T, Youn J-Y, Guttman DS, et al. 2013. The Arabidopsis ZED1 pseudokinase is required for ZAR1-mediated immunity induced by the Pseudomonas syringae type III effector HopZ1a. Proceedings of the National Academy of Sciences 110: 18722–18727.

Lisec J, Meyer RC, Steinfath M, Redestig H, Becher M, Witucka-Wall H, Fiehn O, Törjék O, Selbig J, Altmann T, et al. 2007. Identification of metabolic and biomass QTL in Arabidopsis thaliana in a parallel analysis of RIL and IL populations. The Plant Journal 53: 960–972.

Lisec J, Meyer RC, Steinfath M, Redestig H, Becher M, Witucka-Wall H, Fiehn O, Törjék O, Selbig J, Altmann T, et al. 2008. Identification of metabolic and biomass QTL in Arabidopsis thaliana in a parallel analysis of RIL and IL populations. The Plant Journal 53: 960.

Liu Y, Wu M, Li X, Zhang Y. 2026. Conserved and divergent: Salicylic acid biosynthesis and signaling pathways across the plant kingdom. Molecular Plant 19: 587–605.

Lobell DB, Schlenker W, Costa-Roberts J. 2011. Climate trends and global crop production since 1980. Science 333: 616–620.

Meyer RC, Törjék O, Becher M, Altmann T. 2004. Heterosis of biomass production in Arabidopsis. Establishment during early development. Plant Physiology 134: 1813–1823.

Mishina TE, Zeier J. 2007. Bacterial non-host resistance: Interactions of Arabidopsis with non-adapted Pseudomonas syringae strains. Physiologia Plantarum 131: 448–461.

Mukherjee A, Jodder J, Chowdhury S, Das H, Kundu P. 2025. A novel stress-inducible dCas9 system for solanaceous plants. International Journal of Biological Macromolecules 308: 142462.

Orf I, Tenenboim H, Omranian N, Nikoloski Z, Fernie AR, Lisec J, Brotman Y, Bromke MA. 2022. Transcriptomic and Metabolomic Analysis of a Pseudomonas-Resistant versus a Susceptible Arabidopsis Accession. International Journal of Molecular Sciences 23: 12087.

Rieseberg LH, Archer MA, Wayne RK. 1999. Transgressive segregation, adaptation and speciation. Heredity 83: 363–372.

Sardar A, Nandi AK, Chattopadhyay D. 2017. CBL-interacting protein kinase 6 negatively regulates immune response to Pseudomonas syringae in Arabidopsis. Journal of experimental botany 68: 3573–3584.

Saucet SB, Ma Y, Sarris PF, Furzer OJ, Sohn KH, Jones JDG. 2015. Two linked pairs of Arabidopsis TNL resistance genes independently confer recognition of bacterial effector AvrRps4. Nature Communications 6: 6338.

Savary S, Willocquet L, Pethybridge SJ, Esker P, McRoberts N, Nelson A. 2019. The global burden of pathogens and pests on major food crops. Nature Ecology and Evolution 3: 430–439.

Sun T, Zhang Y, Li Y, Zhang Q, Ding Y, Zhang Y. 2015. ChIP-seq reveals broad roles of SARD1 and CBP60g in regulating plant immunity. Nature Communications 6: 10159.

Törjék O, Berger D, Meyer RC, Mussig C, Schmid KJ, Rosleff Sorensen T, Weisshaar B, Mitchell-Olds T, Altmann T. 2006a. Segregation distortion in Arabidopsis C24/Col-0 and Col-0/C24 recombinant inbred line populations is due to reduced fertility caused by epistatic interaction of two loci. Theoretical and Applied Genetics 113: 1551–1561.

Törjék O, Witucka-Wall H, Meyer RC, von Korff M, Kusterer B, Rautengarten C, Altmann T. 2006b. Segregation distortion in Arabidopsis C24/Col-0 and Col-0/C24 recombinant inbred line populations is due to reduced fertility caused by epistatic interaction of two loci. Theoretical and Applied Genetics 113: 1551–1561.

Velasquez A, Castroverde CDM, He SY. 2018. Pathogen Warfare under Changing Climate Conditions. Current Biology 28: 619–634.

Vishwakarma NK, Yadav S, Sardar A, Choudhary M, Chattopadhyay D. 2026. CBL1/9– CIPK6 complex negatively regulates Respiratory burst oxidase homolog D in Arabidopsis thaliana. The Plant Journal 125: e70700.

Wang Y, Bao Z, Zhu Y, Hua J. 2009a. Analysis of Temperature Modulation of Plant Defense Against Biotrophic Microbes. Molecular Plant-Microbe Interactions® 22: 498–506.

Wang L, Tsuda K, Sato M, Cohen JD, Katagiri F, Glazebrook J. 2009b. Arabidopsis CaM Binding Protein CBP60g Contributes to MAMP-Induced SA Accumulation and Is Involved in Disease Resistance against Pseudomonas syringae. PLOS Pathogens 5: e1000301.

Whitham S, McCormick S, Baker B. 1996. The N gene of tobacco confers resistance to tobacco mosaic virus in transgenic tomato. Proceedings of the National Academy of Sciences 93: 8776–8781.

Zhao C, Liu B, Piao S, Wang X, Lobell DB, Huang Y, Huang M, Yao Y, Bassu S, Ciais P, et al. 2017. Temperature increase reduces global yields of major crops in four independent estimates. Proceedings of the National Academy of Sciences of the United States of America 114: 9326–9331.

Zhu Y, Qian W, Hua J. 2010. Temperature Modulates Plant Defense Responses through NB-LRR Proteins. PLOS Pathogens 6: e1000844.

