## Supplemental Information for "Natural variation in temperature-resilient immunity in Arabidopsis"

Supplemental figures S1-S5

Supplemental tables S1-S3

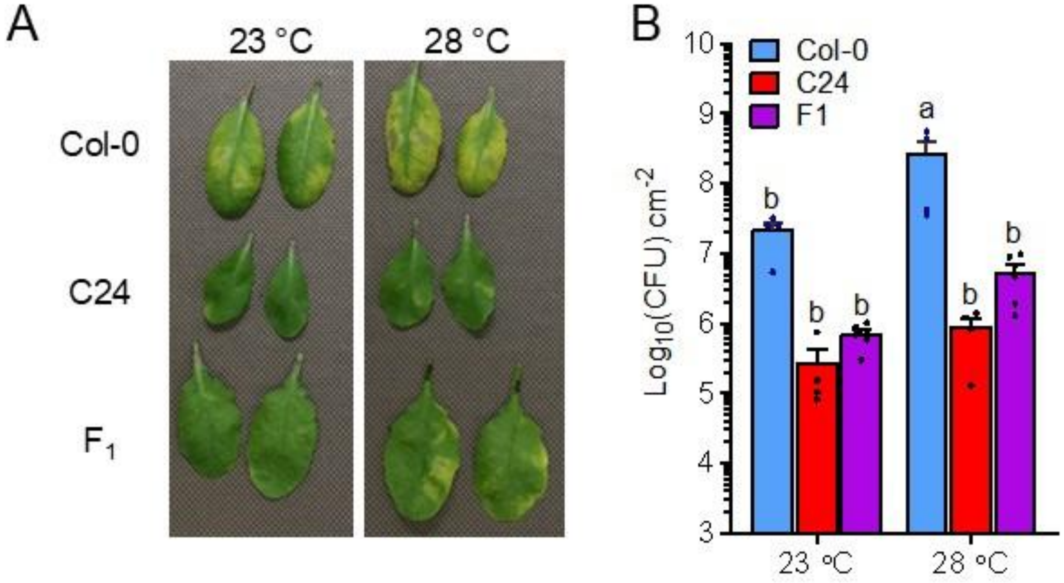


**Figure S1. Col-0 x C24 $F_1$ hybrids exhibit biomass heterosis and disease resistance resilience at elevated temperature. (**A) Images of representative rosette leaves infiltrated with *Pst* DC3000 ($1x10^6$ $CFUml^{-1}$) and (B) bacterial levels at 3 dpi. Data are mean + SD (n=4) and statistical significance was assessed using Two-way ANOVA, with Tukey's honest significance difference (HSD).

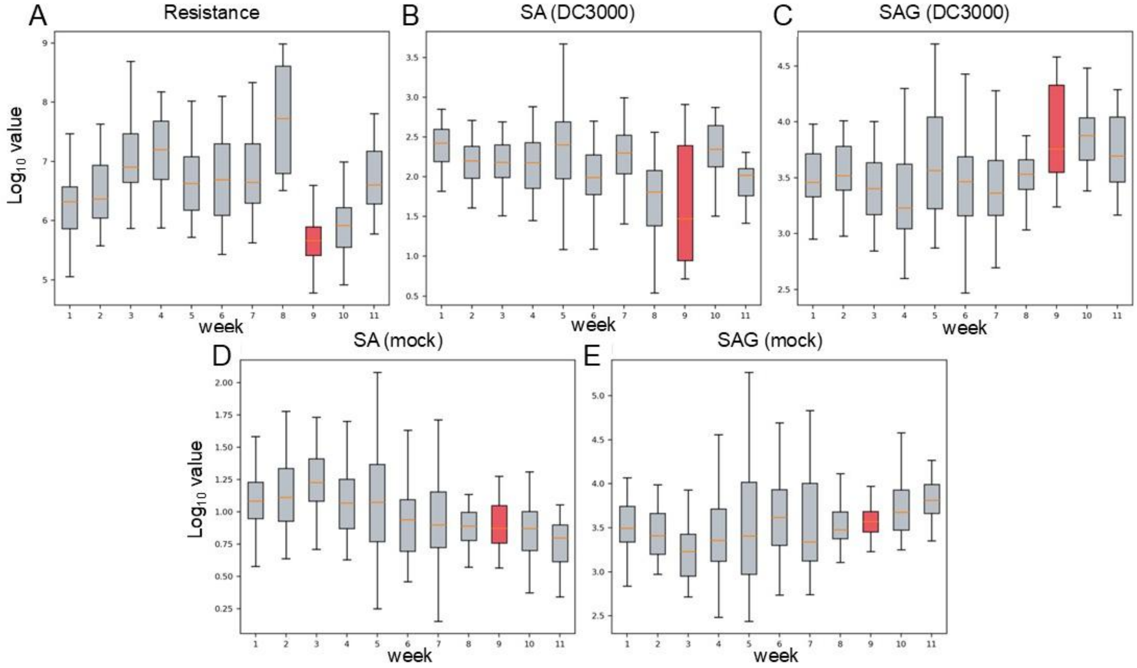


**Figure S2. Phenotyping-batch effects for the five traits in Figure 2.** $Log_{10}$-transformed means for RILs tested per week for (A) Disease, (B) SA (DC3000), (C) SAG (DC3000), (D) SA (mock), and (E) SAG (mock). Each trait is shown by phenotyping batch (weeks 1-11, in assay order). Batch 9 (red boxes) is flagged as a likely failed assay based on its aberrant Col-0 control value (Methods) but is retained in the primary analysis with batch modeled as a covariate.

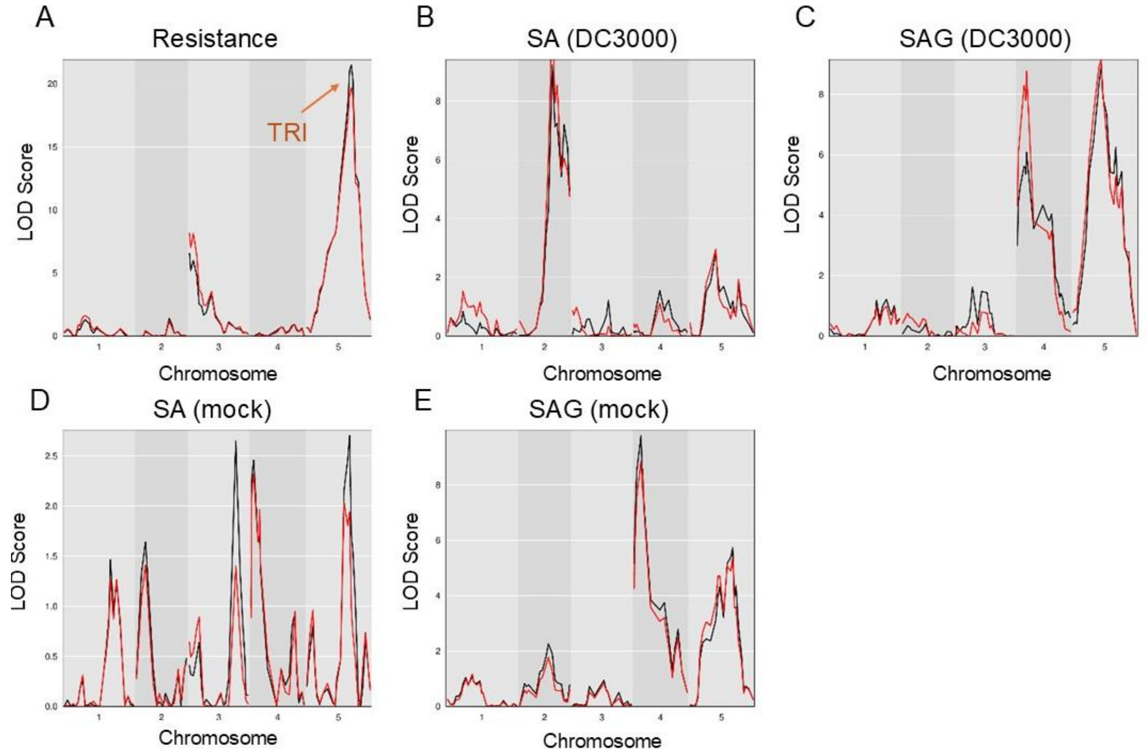


**Figure S3. Effect of Batch-9 exclusion for the genome-wide LOD profiles in Figure 2.** Full dataset (black) overlaid against the analysis with batch 9 excluded dataset (red), for the same five traits and panel order as Figure 2: (A) DC3000 resistance, (B) SA (DC3000), (C) SAG (DC3000), (D) SA (mock), (E) SAG (mock).

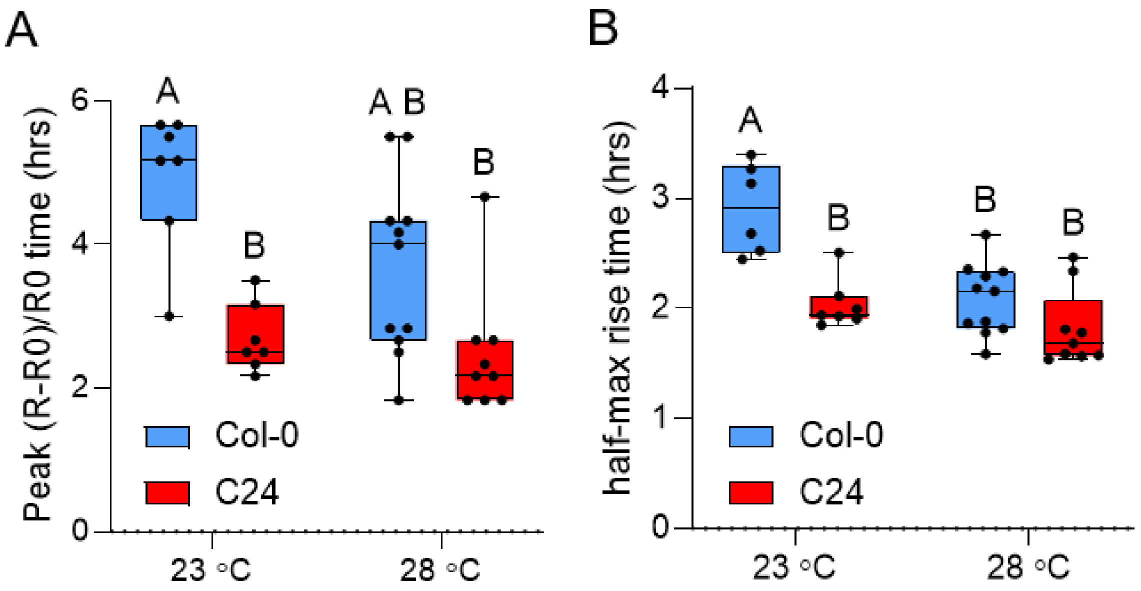


**Figure S4. $Ca^{2+}$ response kinetics during effector-triggered immunity in Col-0 and C24**

The time to reach peak (R-R0)/R0 and the half-maximal rise time for Col-0 and C24 plants infiltrated with *Pst* DC3000 (HopZ1a) from Figure 5D and 5E were assessed. Two-way ANOVA with Tukey's test for significance ($p<0.05$) was performed for peak maxima analyses with letters denoting differences in statistical significance.

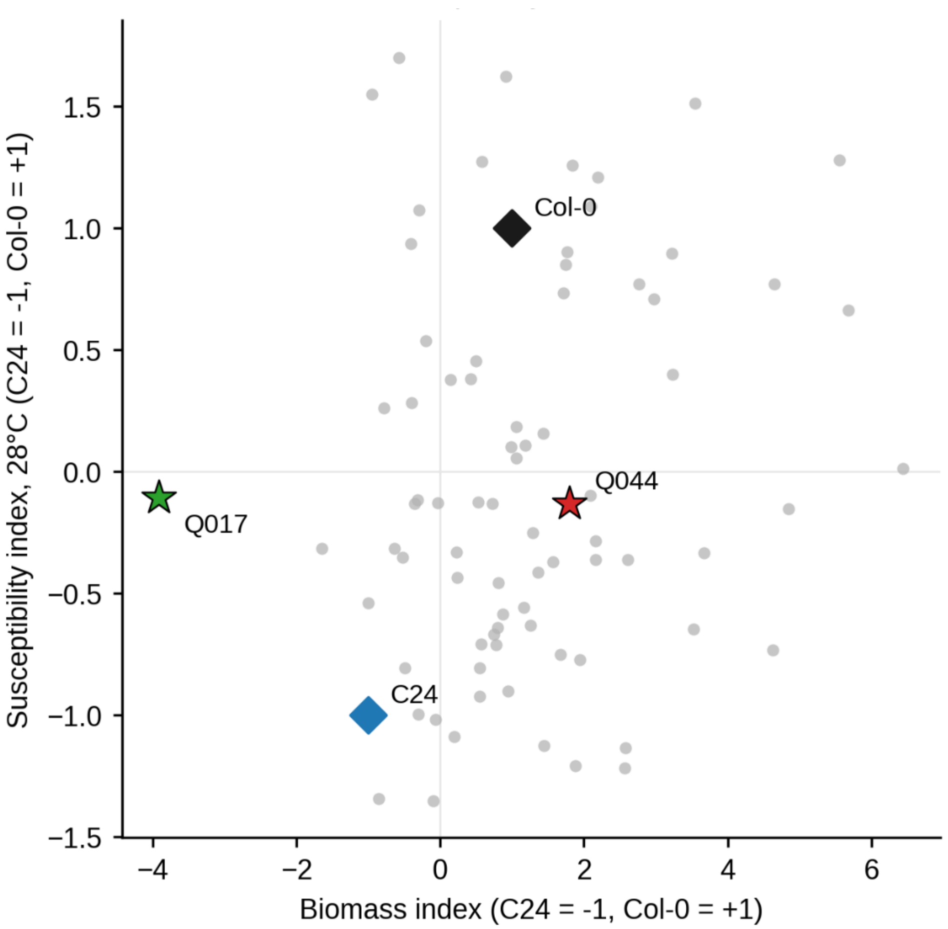


**Figure S5. Biomass vs. disease susceptibility across the dip-screened RIL population at elevated temperature**

Biomass vs. 28°C disease susceptibility across the dip-screen RIL population (gray, n = 77) and Col-0, C24, Q044, Q017 from Figure 6. Axes were normalized so that C24 = −1, Col-0 = +1 (see Methods). RILQ044 and RILQ017 data are from separate experimental batches with Col-0 and C24 Col-0 and C24 as internal controls.

**Table S1. QTL peaks identified in this study.**

| Trait | Chr | Position (cM) | LOD (full dataset) | LOD (no week 9) | 1.5-LOD CI (cM) | PVE (%) | Higher-value allele |
|---|---|---|---|---|---|---|---|
| Disease susceptibility | 3 | 0.0 | 6.57 | 8.16 | 0.0-11.0 | 6.6 | Col-0 |
| Disease susceptibility | 3 | 32.6 | 3.37 | (merged, above) | 20.9-38.2 | 3.5 | Col-0 |
| Disease susceptibility | 5 | 65.5 | 21.50 | 19.67 | 60.1-71.6 | 20.1 | Col-0 |
| SA (DC3000) | 2 | 48.4 | 9.23 | 11.21 | 42.6-52.3 | 9.2 | Col-0 |
| SA (DC3000) | 2 | 65.7 | 7.20 | (merged, above) | 61.8-73.9 | 7.2 | Col-0 |
| SA (DC3000) | 5 | 38.0 | 2.84 | 2.97 | 18.6-76.0 | 2.9 | C24 |
| SAG (DC3000) | 4 | 13.7 | 6.09 | 8.77 | 1.4-24.2 | 6.2 | C24 |
| SAG (DC3000) | 5 | 41.4 | 8.97 | 9.15 | 30.2-49.5 | 8.9 | C24 |
| SAG (mock) | 4 | 10.0 | 9.79 | 8.87 | 1.4-13.7 | 9.8 | C24 |
| SAG (mock) | 5 | 63.0 | 5.74 | 5.40 | 41.7-71.6 / 38.0-65.5* | 5.8 | C24 |

QTL peaks at permutation-based α = 0.05, from the full dataset and the batch-9-excluded dataset. PVE was approximated from the full-analysis LOD score and sample size and was not separately recalculated for the batch-9-excluded dataset. "Higher-value allele" indicates which parental allele is associated with the higher trait value at each peak; for the "disease susceptibility" trait, this analysis identifies the more susceptible allele in Col-0 (i.e., C24 being the more resistant allele) at all three "disease susceptibility" loci), while for the SA/SAG traits it identifies the allele associated with higher defense hormone accumulation, without an implied resistant/susceptible interpretation. *mock_SAG chromosome 5: the 1.5-LOD support interval differs between the full dataset analysis (41.7-71.6 cM) and the batch-9-excluded dataset analysis (38.0-65.5 cM); both are shown because, unlike every other peak in this table, the two analyses gave different results due to batch 9 effect.

**Table S2. Major candidate genes at the chromosome 5 “*TRI*” locus.**

| Gene | Estimated distance to peak | C24 status |
|---|---|---|
| CBL9 (AT5G47100) | 31 kb | Ortholog present |
| VICTR (AT5G46520) | 292 kb | Ortholog present; highly divergent |
| VICTL (AT5G46510) | 298 kb | Ortholog present; highly divergent |
| RPS6 (AT5G46470) | 316 kb | Absent from C24 assembly |
| RRS1B (AT5G45050) / RPS4B (AT5G45060) | ≈980 kb | Ortholog present; highly divergent |
| 4 C24-specific NB-ARC genes (ATC24-5G63040, -63050, -63390, -63480) | ≈800–850 kb* | No Col-0 ortholog |
| 2 C24-specific RSG2-type paralogs (ATC24-5G63330, -63350) | ≈1.6 Mb* | No Col-0 ortholog |
| RLP56 (AT5G49290) | 818 kb | Ortholog present, highly divergent |
| NPR3 (AT5G45110) | 930 kb | Ortholog present, divergent promoter |
| BGLU13 (AT5G44640) | 1.15 Mb | Ortholog present |
| CIPK19 (AT5G45810) / CIPK20 (AT5G45820) | 573–575 kb | Ortholog present |
| AT5G47229 (uncharacterized) | 18 kb | No C24 ortholog |
| AT5G47260 (uncharacterized NLR) | 27.6 kb | Ortholog present |
| ADR1-L3 (AT5G47280) | 31.3 kb | Ortholog present |

Candidate genes were ranked by physical distance to the QTL peak (Chr5:19,161,824; Col-0 genome as reference). *Distance for the C24-specific cluster is measured at its syntenic Col-0 position, since the genes themselves have no Col-0 coordinate. See Dataset S1–S2 for the complete underlying data.

**Table S3. Oligonucleotide primers used for qPCR.**

| Target Gene | Sequences (5'→3') | Reference |
|---|---|---|
| *PP2AA3* (At1G13320) | Forward- GGTTACAAGACAAGGTTCACTC<br>Reverse-CATTCAGGACCAAACTCTTCAG | Huot, *et al.* (2017) |
| *CBP60g* (At5G26920) | Forward - TCGTGGACGCCACCACAAACA<br>Reverse - TCAGCGTTCAGCGGCACGAG | Kim and Castroverde, *et al.* (2022) |

**Supplemental Datasets**

**Datasets S1-11 (separate file).** Candidate genes identified in each QTL peak.

**Dataset S1.** Chr5 "*TRI*" -locus candidates, full and unfiltered (1,339 genes) in the QTL interval Chr5:17,188,344-21,434,621 (TAIR10). Per-gene coding/promoter SNP, indel, and structural-variant counts vs. the C24 assembly (Rowan et al. 2020), plus OrthoFinder ortholog presence/absence and immune-keyword flags.

**Dataset S2.** The 32 C24-specific genes (no Col-0 ortholog) in the syntenic window mapped from the chr5 disease locus, merged with their NB-ARC/P-loop motif calls and local BLOSUM62 homology best-hits against 24 reviewed Arabidopsis NLR reference proteins.

**Dataset S3.** Candidate genes, chromosome 3 "disease resistance" peak A (0.0 cM, Table S2).

**Dataset S4.** Candidate genes, chromosome 3 "disease resistance" peak B (32.6 cM, Table S2).

**Dataset S5.** Candidate genes, chromosome 2 _SA (DC3000) peak A (48.4 cM, Table S2).

**Dataset S6.** Candidate genes, chromosome 2 SA (DC3000) peak B (65.7 cM, Table S2).

**Dataset S7.** Candidate genes, chromosome 5 SA (DC3000) peak (38.0 cM, Table S2).

**Dataset S8.** Candidate genes, chromosome 4 SAG (DC3000) peak (13.7 cM, Table S2).

**Dataset S9.** Candidate genes, chromosome 5 SAG peak (DC3000) peak (41.4 cM, Table S2).

**Dataset S10**. Candidate genes, chromosome 4 SAG (mock) peak (10.0 cM, Table S2).

**Dataset S11.** Candidate genes, chromosome 5 SAG (mock) peak (63.0 cM, Table S2), full-data 1.5-LOD support interval (41.7-71.6 cM).
